# FAIRly big: A framework for computationally reproducible processing of large-scale data

**DOI:** 10.1101/2021.10.12.464122

**Authors:** Adina S. Wagner, Laura K. Waite, Małgorzata Wierzba, Felix Hoffstaedter, Alexander Q. Waite, Benjamin Poldrack, Simon B. Eickhoff, Michael Hanke

## Abstract

Large-scale datasets present unique opportunities to perform scientific investigations with unprecedented breadth. However, they also pose considerable challenges for the findability, accessibility, interoperability, and reusability (FAIR) of research outcomes due to infrastructure limitations, data usage constraints, or software license restrictions. Here we introduce a DataLad-based, domain-agnostic framework suitable for reproducible data processing in compliance with open science mandates. The framework attempts to minimize platform idiosyncrasies and performance-related complexities. It affords the capture of machine-actionable computational provenance records that can be used to retrace and verify the origins of research outcomes, as well as be re-executed independent of the original computing infrastructure. We demonstrate the framework’s performance using two showcases: one highlighting data sharing and transparency (using the studyforrest.org dataset) and another highlighting scalability (using the largest public brain imaging dataset available: the UK Biobank dataset).

---

The amount of data available to researchers has steadily grown, but over the past decade, a focus on diverse, representative samples has resulted in datasets of unprecedented size. The Wind Integration National Dataset (WIND) Toolkit ^1^, CERN data (opendata.cern.ch), or NASA Earth data (earthdata.nasa.gov) are only some of the prominent examples of large, openly shared datasets across scientific disciplines. This development is accompanied by a growing awareness of the importance to make the data more findable, accessible, interoperable, and reusable (FAIR) ^2^, and increasing availability of research standards and tools that facilitate data sharing and management ^3^.

Though large-scale datasets present unique research opportunities, they also constitute immense challenges. Storage and computational demands strain the capabilities of even well-endowed research institutions’ high-performance compute (HPC) infrastructure — rendering the analysis of these datasets unaffordable using methods common in fields accustomed to smaller datasets (e.g. multiple copies of the data, computationally inefficient processing). With the growing complexity of handling large scale datasets, the trustworthiness of derivative data can be at stake as large-scale computations are more difficult to reproduce, comprehend, and verify. Yet especially in the case of large scale datasets, sharing and reusing data derivatives emerges as the most — or sometimes the only — viable way to extend previous work ^4^. It minimizes duplicate efforts to perform resource-heavy, costly computations that also have considerable environmental impact ^5^, and it can open up research on large data to scholars who do not have access to adequate computational resources. In such contexts, data should thus not only be as FAIR as possible, but also handled in a sustainable manner that places data sharing and reuse as a priority.

The challenges of big data are particularly relevant to the life sciences, such as neuroscience or genetics, where datasets scale to millions of files, hundreds of terabytes ^6,7^, acquired from tens of thousands of participants. Well known examples, such as the Human Connectome Project ^8^, the Adolescent Brain Cognitive Development Study (ABCD) ^9^, or the UK Biobank (UKB) project ^10^, contain diverse data ranging from brain imaging to genetics to clinical and non-clinical measures.

In addition, computational processing of biomedical datasets is rarely fully transparent. Often, datasets contain personal data, which imposes usage constraints and prohibits the open distribution of data. Thus, handling these datasets can only be as open as the responsible use of sensitive data permits. Moreover, common processing pipelines possess considerable analytical flexibility, and many tools commonly used in biomedical research rely on proprietary software ^11^, which cannot be easily shared or accessed by others. This threatens the reproducibility of results ^12^, and their *digital provenance* — information about how tools, data, and actors were involved in the generation of a file — is often incomplete. As data processing results often multiply storage demands, the just-keep-everything data management approach is rendered increasingly prohibitive. This fact further impedes the possibility to retrace and verify the origin and provenance of research outcomes fully and transparently ^13^, and hence limits the trustworthiness of the research process and its outcomes ^2^.

Here, we present a portable, free and open source framework — built on DataLad ^14^ and containerization software ^15^ — to reproducibly process large-scale datasets. It empowers independent consumers to verify or reproduce the results based on *machine-actionable* (i.e., machine-readable, automatically re-executable) records of computational provenance, in an infrastructure-agnostic fashion. The framework capitalizes on established technology, used in conjunction with workflows from software development and workload management. Two use cases demonstrate different framework features and its scalability: 1) an application of a MATLAB-based, containerized, neuroimaging processing pipeline on big data from the UKB project ^16^ (comprising 76 TB in 43 million files under strict usage constraints), and 2) a showcase implementation with openly available processing pipeline and data that illustrates the framework’s potential for transparent sharing and reuse of reproducible derivatives. While one can apply the framework by following the description in this work, a bootstrapping script for each use case is provided that — given input dataset and processing pipeline — performs the necessary setup from scratch.

## Results

The proposed framework employs a range of software tools for data, code, and computation management to apply workflows from software engineering — in particular distributed development — to computational research. Specifically, it orchestrates arbitrary data processing via a lean network of interconnected, but self-sufficient workspaces while optimizing for portability, scalability, and automatic computational reproducibility.

To achieve this, our framework combines a range of open source software tools — distributed version control systems, containerization software, job scheduling tools, and storage solutions with optional encryption and compression — into a sequential workflow. A complete, schematic overview is depicted in Figure 1 and basic DataLad concepts are summarized in Box 1. Three key features of this data management solution are central to the framework:

- Comprehensive data structure to track all elements of digital processing
- Computation in automatically bootstrapped ephemeral workspaces
- Process provenance capture in machine-actionable records

**Figure 1:**
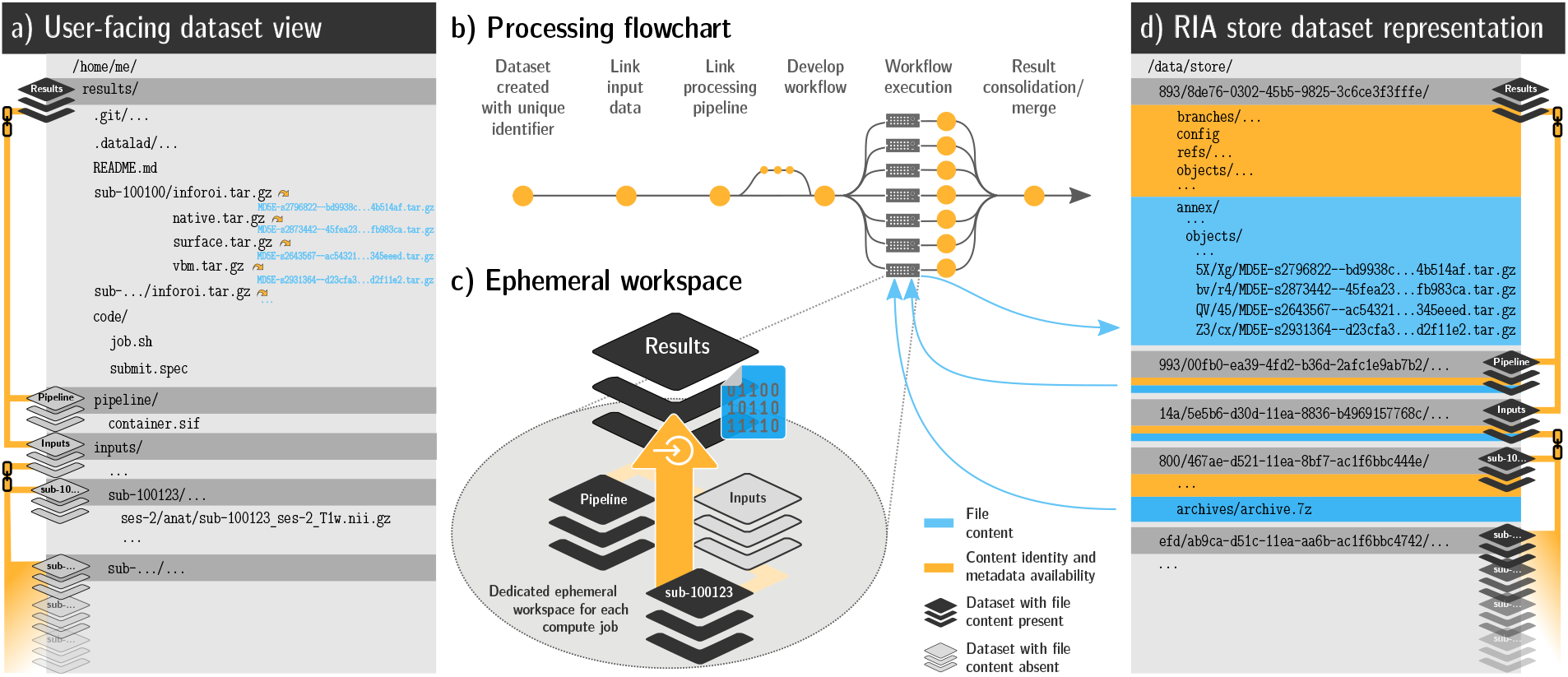
Schematic overview of the processing framework. a) The user-facing representation of the results on a file system after completed processing: A lean DataLad dataset that tracks the computed results, links input data and pipeline, and contains actionable process provenance and location information, allowing on-demand file retrieval or recomputation. Depicted files are from the UK Biobank showcase. b) Process-flowchart: First, a DataLad dataset links required processing components (e.g., input data, processing pipeline, additional scripts). Next, compute jobs are executed, if possible in parallel. Afterwards, results and provenance are aggregated (merged). c) An ephemeral (short-lived) compute workspace: Each compute job creates a temporary, lean *clone*, which *retrieves* only relevant subsets of data, and *captures* the processing execution as provenance. After completion, results and provenance are *pushed* into permanent storage (see d), and the ephemeral workspace is purged. d) The internal dataset representation in a RIA store: The store receives results and can contain input data, optionally using compressed archives (for reduced disk space/inode consumption) or encryption during storage and transport. It is the only place where results take up permanent disk space. If inputs are available from other infrastructure (external, web-accessible servers, cloud infrastructure), jobs can obtain them from registered sources, removing the need for duplicate storage of input data.

#### DataLad concepts

##### DataLad dataset

DataLad’s core data structure is the *dataset*. On a technical level, it is a joint Git/git-annex ^17^ repository. Conceptually, it is an overlay data structure that is particularly suited to address data integration challenges. It enables users to version control files of any size or type, track and transport files in a distributed network of dataset *clones*, as well as record and re-execute actionable process provenance on the genesis of file content. DataLad datasets have the ability to retrieve or drop registered, remote file content on demand with single file granularity. This is possible based on a lean record of file identity and file availability (checksum and URLs) irrespective of the true file size. A user does not need to be aware of the actual download source of a file’s content, as precise file identity is automatically verified regardless of a particular retrieval method, and the specification of redundant sources is supported. These technical features enable the implementation of infrastructure-agnostic data retrieval and deposition logic in user code.

##### A clone

(Git concept) is a copy of a DataLad dataset that is linked to its origin dataset and its history. The clones are lightweight and can typically be obtained within seconds, as they are primarily comprised of file identity and availability records. DataLad enables synchronization of content between clones and, hence, the propagation of updates.

##### A branch

(Git concept) is an independent segment of a DataLad dataset’s history. It enables the separation of parallel developments based on a common starting point. Branches can encompass arbitrarily different modifications of a dataset. In a typical collaborative development or parallel processing routine, changes are initially introduced in branches and are later consolidated by merging them into a mainline branch.

##### Nesting

A DataLad dataset can also contain other DataLad datasets. Analog to file content, this linkage is implemented using a lightweight dataset identity and availability record (based on Git’s submodules). This nesting enables flexible (re-)use of datasets in a different context. For example, it allows for the composition of a project directory from precisely versioned, modular units that unambiguously link all inputs of a project to its outcomes. Nesting offers actionable dataset linkage at virtually no disk space cost, while providing the same on-demand retrieval and deposition convenience as for file content operations because DataLad can work with a hierarchy of nested datasets as if they are a single monolithic repository. When a DataLad dataset B is nested inside DataLad dataset A, we also refer to A as the *superdataset* and to B as a *subdataset*. A superdataset can link any number of subdatasets, and datasets can simultaneously be both super- and subdataset.

##### RIA store

A file-system based store for DataLad datasets with minimal server-side software requirements (in particular no DataLad, no git-annex, and Git only for specific optional features) ^18^. These stores offer inode minimization (using indexed 7-zip archives). A dataset of arbitrary size and number of files can be hosted while consuming fewer than 25 inodes, while nevertheless offering random read access to individual files at a low and constant latency independent of the actual archive size. Combined with optional file content encryption and compression, RIA (“Remote Indexed Archive”) stores are particularly suited for staging large-scale, sensitive data to process on HPC resources.

##### DataLad extension

The core DataLad software is extensible via independently developed Python packages. We developed a custom extension, `datalad-ukbiobank` ^19^ (docs.datalad.org/projects/ukbiobank), to use the UK Biobank (UKB) as a data source for reproducible research. This extension equips Data-Lad with a set of commands to obtain, monitor, and restructure the UKB imaging data release. UKB data are tracked in DataLad datasets that can be updated whenever the UKB updates or adjusts its offerings. Using a multi-branch approach, the DataLad datasets provide a BIDS-structured representation in addition to the UKB-native data organization, without storage duplication and with full provenance capture of the BIDS transformation. We also employed the `datalad-container` ^20^ extension, which integrates container-based command execution with DataLad’s process provenance capture capabilities (see docs.datalad.org/projects/container for more information).

**Box 1:** Main concepts about the design and function of the framework, DataLad, and its underlying technical components. DataLad, integral to the processing framework, is a domain agnostic data management solution based on Git (git-scm.com) and git-annex ^17^. It provides standard interfaces for arbitrary data transport methods, comprehensive process provenance capture for computational reproducibility, and the means to apply proven workflows from collaborative software development to the domain of data processing. More information on DataLad is available at datalad.org and handbook.datalad.org ^21^.

### Comprehensive data structure to track all elements of digital processing

All files involved in processing are contained in DataLad datasets, a Git-repository-based data representation that streamlines data management, sharing, and reuse ^22^. In our framework, such datasets have a common representation (a regular directory tree) familiar to users, and also have a storage representation (a RIA store) that facilitates programmatic data management and reduces storage demands (Figure 1a, d).

DataLad datasets can version control files regardless of file size, and can link other DataLad datasets at precise versions in modular *superdataset-subdataset* relationships. Based on this feature, all processing components, such as data, code, and computational environments in the form of software container images, can be uniquely and transparently identified with single file granularity across a hierarchy of linked DataLad datasets. Unlike purely Git-based tracking, version control and file identification are based on a cryptographic hash of the file content, a feature provided by the software git-annex ^17^. More precisely, each file’s content is translated into a checksum, and this checksum is saved (*committed*) as a file content identifier into the *revision history*—a detailed record of all changes in a DataLad dataset, including their date, time, and author. Exemplary shortened identifiers can be found in Figure 1. This checksum is irreversible, i.e., one cannot infer the file content based on the identifier, but one can verify the content of files that are present on disk. Because file content is not stored in the revision history, the potential to leak sensitive information is significantly reduced, while the data representation still allows for thorough tracking and content verification.

### Computation in automatically bootstrapped ephemeral workspaces

DataLad datasets can be distributed across local or remote infrastructure as lightweight, linked clones. They share their origin dataset’s revision history and can extend it. File content transport across this network is possible via versatile transport logistics that allow for local or remote data hosting. This can enable data transports on systems with too little available disk space for multiple copies, allow redundant storage to be configured, interoperate with hosting services to publish results, or reconfigure data access when remote hosting locations change—without needing to alter the data representation in the dataset.

With these technical features, how and where data are stored (e.g., local, encrypted storage; remote, cloud-based hosting) becomes orthogonal to how and where computations are performed (e.g., on-site compute cluster; remote cloud-computing service). This allows our framework to bootstrap *ephemeral* (short-lived) workspaces for individual computational jobs, retrieve only relevant processing elements (e.g., subsets of input data), and extend the DataLad datasets’ revision history with their results and process provenance (Figure 1c). This, in turn, opens the possibility for parallel *and* version controlled analysis progression, using a distributed network of temporary clones. Results and revision histories can be merged to form a full processing history, in a similar way to how code is collaboratively developed with distributed version control tools ^23^. Importantly, DataLad itself is not a workflow engine, but can be employed for individual nodes and segments of a processing graph defined by other solutions like HTCondor DAGMan ^24^, or snakemake ^25^.

### Process provenance capture in machine-actionable records

Process provenance—how code and commands created results from input data in a particular computational environment—of any processing routine can be captured and stored in machine-readable, automatically re-executable records (Figure 2). These records are created by a datalad run command for the execution of a shell command, or a container invocation by `datalad containers-run`. Users need to supply the command, a software environment, input data, and optionally which results should be saved as parameters. DataLad’s execution wrappers retrieve inputs, initiate command execution, and save results together with a provenance record. Through the use of ephemeral workspaces during provenance capture, the validity and completeness of provenance records is automatically tested: only declared inputs are retrieved, only declared outputs are saved and deposited on permanent storage.

**Figure 2:**
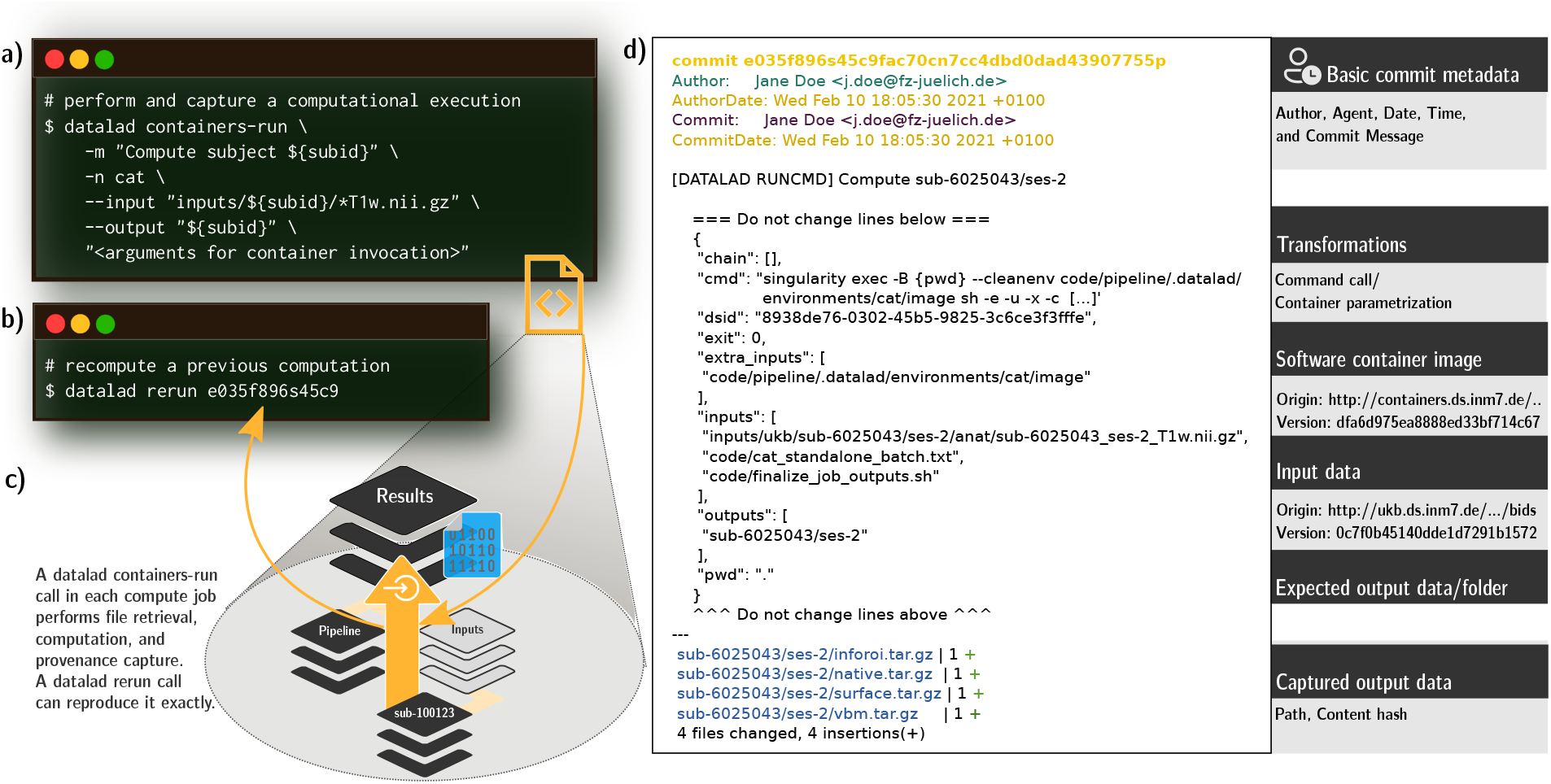
Process provenance of an individual job, its generation, and re-execution. a) Actionable process provenance is generated with a datalad containers-run command. This example contains a container name specification (`cat`), a container parametrization or command, a commit message, and an input and output data specification. The provenance is stored as a structured, JSON-formatted record linked to a Git commit. b) To re-execute a process, the `datalad rerun` command only needs to be parameterized with a revision identifier, such as a Git tag, a “commit shasum” (`e035f896s45c9fa[...]` in this example), or a revision range containing one or more commits with associated provenance records. c) The `datalad containers-run` call is at the center of each individual job. As the core execution command (see Listing 1, line 33-39), it performs data retrieval, container execution, and result capture, and generates the actionable provenance that a subsequent `datalad rerun` command (b) can re-execute. With complete provenance, a re-execution is supported on the original hardware, or on different infrastructure. d) The machine-readable, re-executable provenance record stored alongside computed results in the revision history. A legend (right) highlights the most important pieces of recorded provenance. While *automatic* re-execution requires the tool DataLad, sufficient information to repeat a computation using other means can also be inferred from the structured JSON records by other software or even humans. This information forms the basis for standardized provenance reporting, for example using the PROV data model ^27^.

A resulting process provenance record is identified with one unique, hash-based identifier in the revision history, and can subsequently be used by authorized actors to automatically retrieve required components and re-execute the processing, irrespective of whether the original compute infrastructure is available ^26^. This potential for full computational reproducibility of arbitrary processing steps not only increases the trustworthiness of the research process per se, but permits structured investigations of result variability, and furthermore provides the means to rerun any analysis on new data or a updated analysis components.

### Showcases

These DataLad features offer great flexibility for transparently conducting reproducible, high-performance data processing in a wide variety of computational environments. In two concrete showcases we next highlight 1) the scalability of this approach, and 2) complete transparency and reproducibility of this data processing method, when combined with open data and open source tools.

### Use case: large-scale medical imaging data processing

To demonstrate the framework’s scalability, we conducted containerized analyses on one of the largest brain imaging datasets, the UKB imaging data. The strain that this dataset places on computational hardware is considerable both in terms of disk space usage (i.e., the amount of data that a hard drive can store) and inode usage (i.e., the number of files that a file system can index). To show how the framework can mitigate hardware limitations, we processed the dataset on two different infrastructures, an HPC system with inode constraints that preclude storage of the full number of files, and a high throughput computing (HTC) system with disk space limitations that preclude data duplication. In doing so, we assessed if the framework can be used across different infrastructures, investigated result variability between two recomputations of the pipeline, and probed the framework’s features under distribution restrictions of both the data and the MATLAB-based software component. Finally, in order to demonstrate that the framework can capture and re-execute complete process provenance, we also recomputed individual results on a personal laptop.

As a first step, we prepared input data and computational pipeline. We created a Singularity container with a pipeline to perform voxel-based morphometry (VBM) ^28^ on individual T1-weighted MRI images based on the Computational Anatomy Toolbox (CAT) ^29^. We stored UKB data in archives in a DataLad RIA store ^18^ (Figure 1d) to mitigate disk-space and inode limitations on the two different systems. The store comprised 42,715 BIDS-structured ^30^ DataLad datasets, one per study participant, that were jointly tracked by a single additional superdataset (UKB-BIDS; “Data” in Figure 3). In total, the domain-agnostic data representation in the store hosted 76 TB of version-controlled data with 43 million individually accessible files while consuming less than 940k inodes.

**Figure 3:**
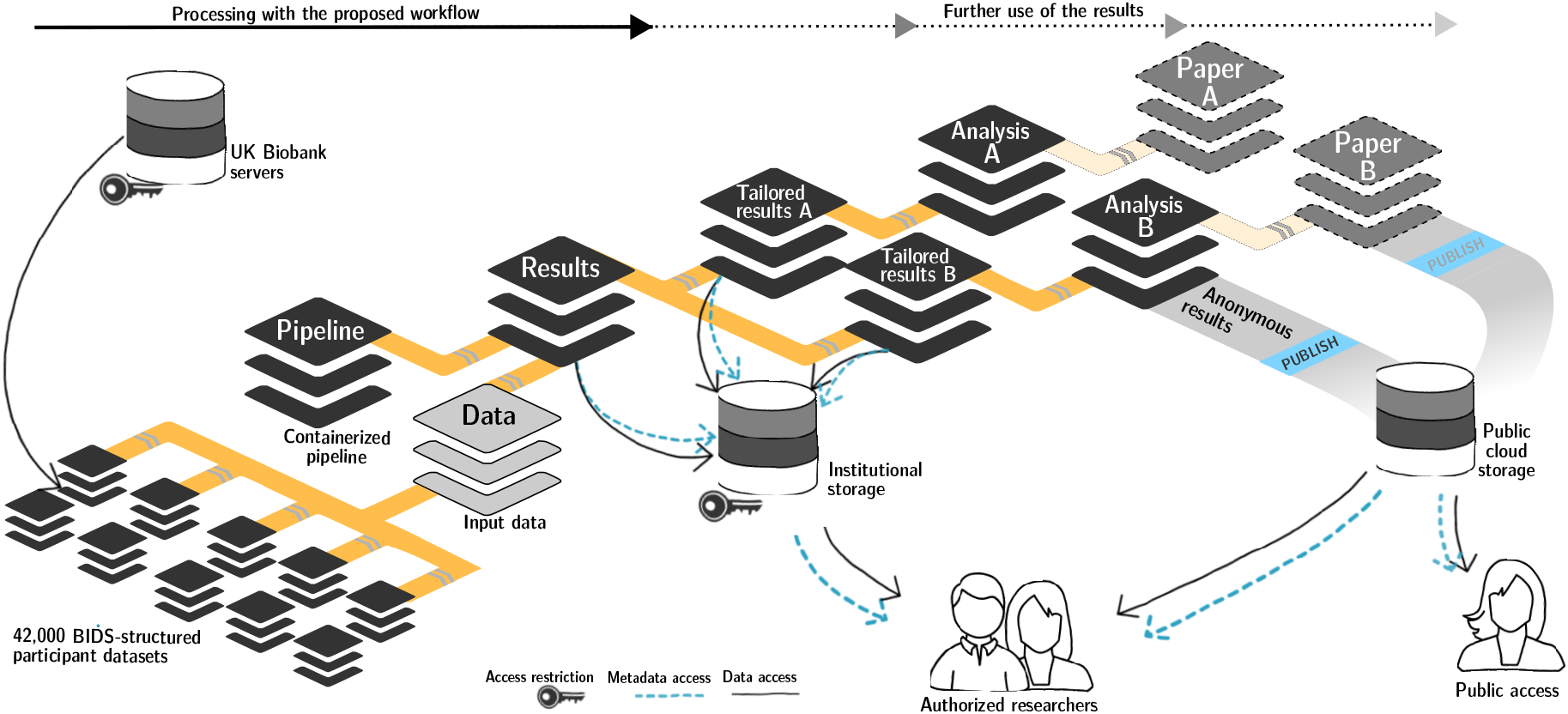
Overview of DataLad dataset linkage through processing and reuse. Any DataLad dataset may comprise other DataLad datasets as *subdatasets* via lightweight but actionable and versioned links. This connects a dataset to the content and provenance of a different modular unit of data, such as the outcomes of a the preceding processing step. The genesis of an analysis output (*Analysis A/B*) based on intermediate processing outcomes (*Tailored results A/B*) can thus be traced back all the way to the original raw data. Access control and storage choices are independent across individual components in this network of linked data modules. Aggregated data and analysis results can be shared with larger audiences or publicly on a variety of platforms, while raw and derived data may underlie particular access restrictions, or require particular hosting solutions due to their size.

Next, we assembled a single DataLad dataset to capture all processing inputs and outputs (“Results” in Figure 3). It initially tracked: 1) the UKB-BIDS DataLad dataset; 2) a DataLad dataset providing the containerized CAT pipeline; 3) the compute job implementation responsible for bootstrapping a temporary workspace, performing a parameterized execution of the pipeline, capturing its outputs, and depositing results in the RIA store (see Listing 1 for a simplified version); and 4) the job scheduling specifications for SLURM ^31^ (used on the HPC system) and HTCondor ^24^ (used on the HTC system). Despite the total size of all tracked components, the pre-execution state of this dataset was extremely lean, as only availability (a URL) and joint identity (single checksum) information on the linked datasets is stored, and all other information is contained in the linked datasets themselves. This also implies that the DataLad dataset tracking the computational outputs is *not* automatically encumbered with sensitive information, even though it precisely identifies the medical imaging input data.

The compute job implementation minimized the number of output files using tar archives to reduce the strain on the technical infrastructure, and removed undesired sources of result variability (time stamps, file order differences in archives, etc.) to allow comparisons between recomputations. Later, these archives were partially extracted into tailored result datasets for easy consumption (see Methods, “(Re)use”). To maximize practical reproducibility of computational outcomes, a compute job implementation does not reference any system-specifics, such as absolute paths, or programs and services not tracked and provided by the DataLad dataset itself. This means that any system with DataLad installed, the ability to execute Singularity container images, and a basic UNIX shell environment is capable of recomputing captured outputs. Any performance-related adaptations to the particular systems used for our computations were strictly limited to the job scheduling layer, which is clearly separated from the processing pipeline. Computation and recomputation on systems with different batch scheduling software is then possible by providing alternative job specifications, without changes to the pipeline implementation.

We performed processing on the HPC and HTC infrastructure starting from the exact same dataset version state, but with job orchestration tuned to the respective job scheduling system^1^. Provenance for each execution of the CAT pipeline on an individual image was captured in a dedicated commit, and recorded on a participant-specific Git branch. Recorded outputs and provenance records were pushed to the RIA store on job completion, yielding a total of 995.6 GB of computed derivatives in 163,212 files. The second computation added matching commits and branches to the DataLad dataset that enabled straightforward comparison and visualization of results using standard Git tools and workflows. To confirm the practicality of computational reproducibility solely based on the captured computational provenance information, we performed automatic recomputation of individual results on a consumer-grade, personal laptop without job scheduling. This type of spot-checking results resembles the scenario of an interested reader or reviewer of a scientific publication with access to (parts of) the data, but no access to adequate large-scale computing resources.

With the exception of execution time, the number of jobs, proportion of successful jobs, and size and structure of the results were identical between the two systems. Specifically, with the exception of one output flavor (projections of computed estimates to the cortical surface) more than 50% of all output files were identical across the two computations. Outcome variability for non-identical results was largely attributable to minor numerical differences, as illustrated by the mean squared error (MSE) over recomputations for a range of key VBM estimates: total surface area (*μ* = 1891, *MSE* = 0.315), cerebro-spinal fluid (*μ* = 365, *MSE* = 0.052), total intracranial volume (*μ* = 1508, *MSE* = 0.052), white matter (*μ* = 519, *MSE* = 0.004), and gray matter (*μ* = 621, *MSE* = 0.001). We also computed correlations over different brain parcellations included in the CAT toolbox. The lowest observed correlation across recomputations for VBM estimate distribution across different brain parcellations were Pearson’s *ρ* > 0.998 for the Destrieux 2009 surface parcellation ^32^ for all brain regions. Quality control metrics depicted in Figure 4 exhibit *ρ* > 0.99999 for computation and recomputation.

**Figure 4:**
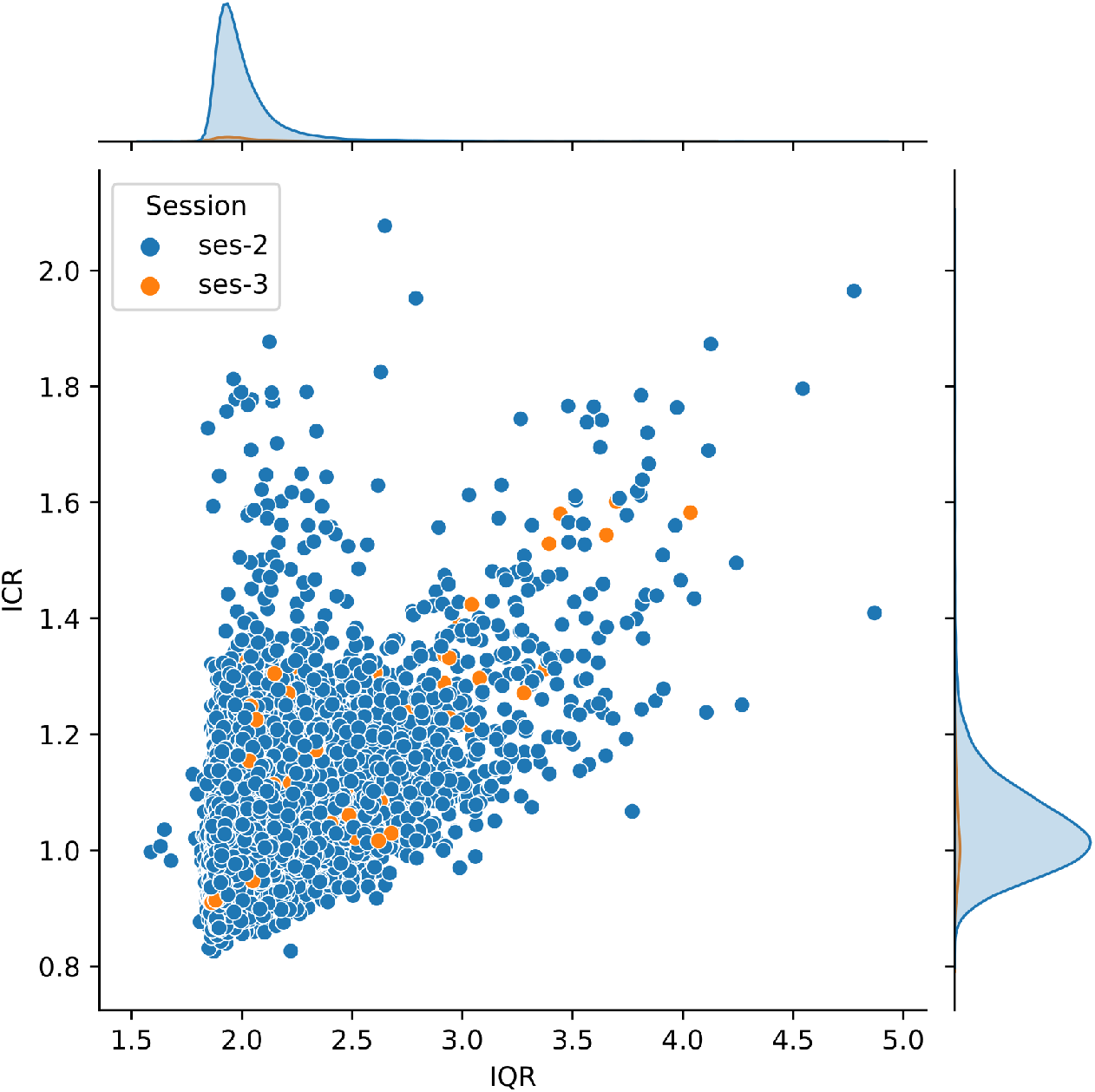
Distribution of quality assurance measures, derived for 41,180 unprocessed T1-weighted images from the UKB dataset. The quality measures were obtained retrospectively based on the preprocessing methods ^47,48^. For both measures, lower values correspond to better quality. Abbreviations: IQR - image quality rating, a weighted composite score based on noise, inhomogeneity, and image resolution (0.5-1.5 = “perfect”, 1.5-2.5 = “good”, 2.5-3.5 = “average”, 3.5-4.5 = “poor”, 4.5-5.5 = “critical”, >5.5 = “unacceptable”); ICR - inhomogeneity contrast ratio, estimated as the standard deviation within the white matter segment of the intensity scaled image.

The complete implementation of this showcase cannot be shared due to imposed data usage and software license restrictions. However, we provide a bootstrapping script that implements all required setup steps at github.com/psychoinformatics-de/fairly-big-processing-workflow, and share a detailed description and full recipe of the container together with instructions on how to build and use it at github.com/m-wierzba/cat-container.

### Use case: Open tutorial

As strict software license restrictions and data usage agreements prevent fully open sharing of computed results and a public demonstration of their process provenance records, we set up an open tutorial analysis using free and open source `fmriprep` software ^33^ and open data from the studyforrest.org project ^34^. We confirmed that process provenance was sufficient to enable automatic recomputations on an HTC system, a personal work station running Debian, and a Mac, and published the resulting DataLad dataset to public GitHub (github.com/psychoinformatics-de/fairly-big-processing-workflow-tutorial) and Gin (gin.g-node.org/adswa/processing-workflow-tutorial) repositories. This demonstrator allows for in-depth inspection, retrieval (`datalad get`) of any and all data processing inputs and outputs, as well as automatic recomputation (`datalad rerun`) of all captured results.

## Discussion

The proposed framework aims to make the results of any processing as open and reusable as the given limits of individual components allow. It streamlines computation, re-computation, and sharing with appropriate audiences for datasets and on compute infrastructure of any size. To this end, proven procedures from software development and a set of open source software tools are assembled into a scalable and portable framework with a variety of features: The basis for transparency is laid with version control for all involved files, including software environments. Distributed data transport and storage logistics offer flexibility to adapt to particular computing infrastructure. Reproducible results are enabled via comprehensive capture of machine-actionable process provenance records, capitalizing on portable containerized environments. Combining distributed computing with ephemeral workspaces that resemble workflows from collaborative software development yields efficient processing, and ensures the validity of provenance information.

The framework shares features and goals with a number of related systems, some of which we want to highlight in order to illustrate how the proposed workflow and its main building blocks relate to other solutions. The proposed *Pan-Neuro*^35^ is an alternative solution for neuroscientific computing on large-scale data, but is more geared towards interactive processing, and represents a centralized, cloud-based platform for computing and data hosting. *IPFS*, the InterPlanetary File System (https://ipfs.io), is a distributed system for data transport that employs an approach to content addressing that is based on cryptographic hashes of file content. This concept is identical to the one employed by git-annex, except for differences in the hash or key composition details. Consequently, these systems are interoperable, and the proposed framework could directly employ IPFS-based data sources via the git-annex integration (https://git-annex.branchable.com/special_remotes/ipfs/). *DVC* ^36^ is a version control system and workflow manager also built on Git. It employs distributed version control for individual large files or collections of files, and captures provenance of language-agnostic machine-learning pipelines that connect multiple steps of building an ML model into a directed acyclic graph (DAG). Major differences lie in DVC’s specific focus on machine learning models in workflow management, and less emphasis on portability and reproducible environments. *Snakemake* ^25^ is a feature-rich and domain-agnostic workflow engine, with support for including software environments in the form of conda-environments or software containers, portable workflows that allow execution on remote resources such as cloud services or batch systems on compute clusters, and provenance capture. DataLad can enhance snakemake workflows by retrieving versioned input data files (http://docs.datalad.org/projects/mihextras/generated/man/datalad-x-snakemake.html), and snakemake-based compute job orchestration could be employed as an alternative to the custom implementations for SLURM and HTCondor used in the work presented here. *Parsl* ^37^ is a Python-based parallel scripting library that is designed to enable compositional programming for a variety of scientific use cases. It features workflow management, a mix-and-match style portability with provider interfaces to configure resource-specific requirements across tools or infrastructures, and data management that can perform data transfers to and from resources via several protocols. Similar to snakemake, it provides an alternative workflow definition and orchestration solution, and could employ the provenance capture approach proposed here. Unlike the framework proposed here, Parsl does not build on a version-controlled core data structure, but on a specification of interconnected apps and the data flow between them. However, like snakemake files, these specifications could be tracked within DataLad datasets in order to combine the capabilities of these systems. *Apache Spark* ^38^ and *Dask* ^39^ are both feature-rich, distributed computing solutions targeting different software ecosystems. Unlike the proposed framework or the aforementioned solutions they require the deployment of dedicated services across a distributed computing resource, and can perform large-scale computations via internal parallelization, where a comparable compute-node local provenance capture step is not performed or possibly even meaningful.

Overall, our framework is a general-purpose solution that is compatible with any data that can be represented as files of any size, and any computation that can be performed via a command line call. It is built on a collection of portable, free and open-source tools that can be deployed without special privileges or administrative coordination on standard HTC/HPC infrastructure, or personal computing equipment.

While it is an explicit aim for the framework to yield FAIR outputs, this aspirational goal is not fully reached. Metadata used and produced by the framework does not conform to explicit annotation standards. Instead, it encodes essential metadata, such as author, date, time, and description in locations that are provided by the version control system Git. Other metadata are put into plain-text, key-value data structures that conform to no particular formal ontology or vocabulary. This shortcoming limits the findability and accessibility of its outputs severely. Questions like “which outcomes were computed with a specific version of a particular software?” cannot be reasonably answered without additional standardization and annotation effort.

That being said, the main contribution of the proposed computational approach is the association of process provenance with captured outcomes (FAIR R1.2), with precise linkage of any data inputs within and across individual datasets (FAIR I3), using unique, content-hash based identifiers for all components (FAIR F1). These metadata are tracked in a dedicated overlay data structure that can ensure their accessibility, even when the underlying data are no longer available or a particular entity has no permission to access them (FAIR A2).

What the framework provides today is a technical system that, despite its ignorance regarding formal metadata standards affords practical, automatic recomputation of arbitrary data processing results. This ability dramatically elevates the starting point for future FAIRification efforts of computational outcomes. Reproducibility can be programmatically verified, thereby providing a confirmation of the comprehensiveness of data and essential process metadata encoded in a DataLad dataset. Subsequent annotations of precisely versioned data, or tracked computational environments can retroactivily boost findability and accessibility of outcomes. For example, an added annotation of the composition of an employed containerized pipeline can help answer the question posed above. Neither metadata format nor terminology are constrained by the proposed framework. Importantly, the ability to recompute outcomes provides a strong incentive for researchers to produce computational outcomes with verifiably complete (meta)data. This is an important half-step towards a FAIRer future that boosts the availability of research outputs that can receive continuous updates to co-evolve with further developments of metadata standards and requirements of future metadata-driven applications. To this end, a DataLad dataset can also be exported to different formats used in frameworks with a similar aim, such as `BagIt`^40^.

The use of containerized software environments plays a key role in the proposed framework. They represent the most practical solution to portable computational environments today. However, their long-term, universal accessibility is all but guaranteed. Even today the `singularity` software does not support all major operating systems. Ten years ago, the popular `docker` software did not yet exist, and it is unclear whether its container images will be executable in ten years from now. Providing the build instructions for a container image, rather than (or in addition to) the readily executable image, may improve the longevity of their accessibility, and also mitigate the problem of license-imposed sharing restriction. However, it is not guaranteed that executing a recipe twice results in identical software containers. *Reproducible builds*, the practice of creating identical container images from a recipe ^41^, for example, require the specification and availability of software and system libraries at precise versions. For the same reason of long-term accessibility, it will also be necessary to incorporate DataLad’s own idiosyncratic provenance metadata into such a comprehensive provenance report — then matching a format and standardization desirable for a particular scope or application. A promising effort towards portability and longevity of container technology is the Open Container Initiative (OCI; opencontainers.org), which aims to create open, vendor-neutral, and portable industry standards around containers formats and runtimes.

Based on process provenance and version control, structured analyses of variability between (re)computations on the same or different infrastructure are facilitated ^13,42^. Bit-identical recomputation of a result are trivially verifiable. The comprehensive capture of input data version, computational environments and process parameterization enable deep inspection of other sources of result variability. Building on this foundation, more standardized process descriptions ^43^ and reproducible computational environments ^41^ can further enhance these types of analyses. Nevertheless, computational results that do not reproduce exactly are a challenge for content checksum based version control systems like Git. If irreproducibility is solely caused by issues of numerical precision and reproducibility of basic floating point operations, it may be possible to nevertheless achieve bitwise identical result by reducing the precision of stored outcomes to empirically meaningful levels of detail, like we did here for the aggregated brain structure scores in the UKB use case. However, in general data type specific and research focus specific implementations of “identity” operators would be required, and while Git offers means for their integration (so-called diff drivers), there are no generally applicable off-the-shelf solutions available for this problem.

The approach to reproducible computation proposed here is applicable to a wide range of use cases. Different datasets, different processing pipelines, and different containerization technologies, such as Docker, can be employed by simply replacing the respective components, and utilizing features already built into DataLad. These possibilities are illustrated in the extensive DataLad Handbook ^21^ at handbook.datalad.org. Combining this framework with more capable workflow engines for defining and orchestrating compute jobs and their interdependencies in the future, would open up the possibility for implementing other types of data processing that go beyond the parallel execution of mutually independent compute jobs that make up the use case illustrated in this work. Such an implementation would need to fulfill three general requirements that were implemented here: 1) The execution of a compute job must take place in a workspace that is defined by a recorded state in a DataLad dataset. 2) Execution via Datalad captures process provenance in a new, incremental dataset state. 3) Advanced dataset states after any parallel processing stage are consolidated into a merged mainline. When these conditions are met, the structure of a processing DAG would be reflected in the version-control history of a DataLad dataset hierarchy that comprises all inputs and outputs of a computational project, with each recorded step being associated with machine-actionable provenance records that enable deep inspection large-scale computing outcomes.

The presented showcases provide two concrete examples for the adoption of the proposed framework that deal with typical obstacles for transparent, reproducible science. The UKB’s data size exceeds the capacity of most infrastructure. We demonstrated the scalability of our framework by processing these data on systems with hardware limitations that would typically render even storage of inputs and outputs difficult or impossible. As the proposed framework enables selective recomputation even on commodity hardware, consumers can investigate results without having to rely on the original authors, and without access to the original computational infrastructure. Even though the raw data may be too large to allow users a complete recomputation, the process provenance entails a trail of processing steps that permits automatic recomputation of individual results. A one-time computation on larger infrastructure can thus build a verifiable, trustworthy foundation for numerous subsequent analyses by other researchers.

Finally, over and above everything else, the framework makes research as open as desired. The medical imaging showcase featured a processing pipeline based on proprietary software and pseudonomized personal data under usage constraints. Data and computational environment are not publicly shareable. But if data usage agreements and software licensing permit, as it is the case in the second showcase, processing results can be shared publicly that are independently and automatically reproducible by any interested party. This level of transparency dramatically improves the accessibility of scientific outcomes.

## Methods

The proposed framework aids the reproducible execution of a containerized pipeline on input data, by associating computational outcomes with machine-actionable provenance records in a version control system. We illustrate the technical details of this process with two use cases that differ in scale as well as data access and processing requirements, but follow a common pattern in general setup and composition.

### Framework setup

The technical nature of the framework components, in particular its foundation, the version control software Git, enables distributed computational workflows that utilize and extend established procedures from collaborative software development to data processing. The framework is bootstrapped in two steps that could be performed by a tailored shell script for a particular application.

#### Self-contained processing specification as a DataLad dataset

The first step is the creation of a new DataLad dataset that will eventually track the processing results (dataset labeled “Results” in Figure 3). Input data, images of containerized pipelines, or custom code are added to this dataset. While the use of software containers to provide processing pipelines is not strictly required, they are a practical method to provide stable and portable computational environments. Because such containers can be stored in image files, they can be tracked and precisely versioned like any other component of a DataLad dataset. The `datalad-container` extension provides a convenience interface for registering containers and for executing commands in such environments.

All processing components, such as processing-specific, customized scripts and applications or data, can be added directly to the dataset as individual files. More typically, however, individual processing components, for example input data or containerized pipelines, are placed in separate Data-Lad datasets and linked as subdatasets (Figure 3). This more modular structure enables (re)use of independently maintained components, while strictly separating access modalities to each of them. In this way, access-restricted input data does not impair sharing of less sensitive outcomes and the versioned link between superdataset and subdatasets guarantees precise identification of processing components, regardless of whether a particular dataset consumer has access to a given component.

The resulting dataset is the entry point to a self-contained directory structure, potentially comprising other nested DataLad datasets, that jointly define identity and location of all data processing inputs in the exact form needed for a particular computation.

#### Environment and performance optimized orchestration

The second step is the preparation of the computational environment and processing orchestration. This relates to *what* is computed as well as *how* it is computed. The compute job orchestration, the *how-to-compute*, could be as simple as direct, sequential executions of required processing steps in a shell script for-loop. However, large-scale computations typically require some form of parallelization. The compute job orchestration is thus likely to be implemented using the job scheduling system of a given compute infrastructure. As such, *how-to-compute* is highly infrastructure-specific, and must determine an optimum balance of resource demands, such as run time, memory and storage requirements, in order to achieve optimal throughput.

The *what-to-compute*, the computational instructions, pipelines, or scripts, need to be independent computational units that can be executed in parallel. A common example is the parallelized execution of a processing pipeline on different, independent parts of input data. As parallelization often corresponds to the granularity at which a recomputation will be possible in our framework, relevant considerations are, for example: “What is the smallest unit for which a recomputation is desirable?”, or “For which unit size is a recomputation still feasible on commodity hardware?”. To ensure reproducibility for an audience that does not have access to the original infrastructure, *what-to-compute* needs to be infrastructure-agnostic, without references to system-specifics such as absolute paths, or programs and services not tracked and provided by the DataLad dataset itself. Then, computation and recomputation of *what-to-compute* are possible on different systems, with any potential adjustments only relating to the job orchestration layer in *how-to-compute*.

### Execution and result consolidation workflow

After the two preparatory steps are completed the actual data processing can be executed by submitting the compute jobs to the job scheduling system. Each compute job will clone the DataLad dataset with the processing specification to a temporary location, bootstrap an ephemeral workspace that is populated with all inputs required for the given job with a job-specific parameterization, execute the desired computing pipeline, and capture a precise provenance record of this execution, comprising all inputs, parameters and generated outcomes. Lastly, it pushes this provenance metadata and result file content to permanent storage. This workflow resembles a standard distributed development workflow in software projects (obtain a development snapshot, implement a new feature, and integrate the contribution with the mainline development and other simultaneously executed developments) but applies it to processing of data of any size. Specific details of this workflow are outlined in sequential order in the following paragraphs. Where applicable, they annotate and rationalize the generic compute job implementation in Listing 1.

#### Dataset clone source and update push target

can be separated in an initial setup step to improve performance. When all compute jobs deposit their outcomes at the same DataLad dataset location that later compute jobs also clone from, version history in this dataset accumulates and progressively slows the bootstrapping of work environments of compute jobs, because more information needs to be transferred. Moreover, result deposition in a DataLad dataset is a write operation that must be protected against concurrent read and write access for technical reasons, and hence introduces a throughput bottleneck. Both problems are addressed by placing an additional clone of the pre-computation state of the processing specification dataset in a RIA store before job submission (Figure 1d). This clone is used for result deposition only (Listing 1, lines 17 and 49). Dataset clones performed by jobs are done from the original location that is never updated, hence also never grows. In order to avoid unintentional modifications during long computations, the dataset clone source for jobs may not be the dataset location used for preparation (Figure 1a), but yet another, separate clone in a different RIA store. The clone source and push target locations are provided as parameters to compute jobs (Listing 1, lines 5-6). All dataset locations are not confined to exist on the same hardware as long as they are accessible via supported data transport mechanisms over the network.

#### Job-specific ephemeral workspaces

are the centerpiece of the computation, and the location where the actual data processing takes place (Figure 1c). Critically, these workspaces are boot-strapped using information from the specification DataLad dataset only. This is achieved by cloning this dataset into the workspace first (Listing 1, line 11), and subsequently performing all operations in the context of the clone. After computation and result deposition the clone and the entire workspace are purged. This ensures that all information required to perform a computation is encoded in this portable specification, that it is actionable enough to create a suitable computing environment, and that all desired outcomes are properly registered with the DataLad dataset to achieve deposition on permanent storage.

#### Containerized execution and provenance capture

happens within the ephemeral workspace on a uniquely identified branch per job (Figure 1b, “workflow execution”; Listing 1, line 21). Prior computation, the state of this branch is identical for all jobs. It comprehensively and precisely identifies all processing inputs, and links them to author identities, time stamps, and human readable descriptions encoded in the Git revision history of the dataset (Figure 2d, top).

Based on this initial state, a computational pipeline is then executed, and all relevant computational outcomes are saved to the DataLad dataset to form an updated state (Listing 1, line 33-39). For this execution, all required input files are specified by their relative path in the DataLad dataset (potentially pointing into linked subdatasets). Importantly, only these job-specific inputs will be transferred to the compute job’s environment. Likewise to be saved outcomes are selected by providing path specifications. Given the execution of the computation in an isolated, ephemeral workspace that is unique for each individual job, two guarantees can be derived regarding the provenance of the computational outcomes: 1) All dataset modifications can be causally attributed to the initiated computation; 2) only declared inputs were required to produce the outcomes.

DataLad commands like `run` (for command line execution) or `containers-run` (for execution in containerized environments) yield machine-readable provenance records that express what command was executed, with which exact parameters, based on which inputs, to generate a set of output files (Figure 2d). Such a record is embedded in the Git commit message of the newly saved dataset state as structured data. The record itself is lean and free off explicit version information for individual inputs, because the dataset state as a whole jointly identifies all versions of all dataset components, such that individual versions are readily retrievable on a (later) inspection of this state.

The captured provenance record is machine-actionable. Using the dataset and information in the provenance record in a dataset state’s commit message, the DataLad command `rerun` can reobtain necessary inputs and run the exact same command again, given availability of data and environments. Importantly, this re-execution does not strictly depend on the original compute infrastructure, but benefits from DataLad’s ability to retrieve file content from multiple redundant locations.

#### Result deposition

takes place after successful completion of each job. The file content of computational outcomes, along with their provenance, are pushed to permanent storage (Figure 1b, blue arrow). Two different components of result deposition have to be distinguished.

Transfer of file content (Listing 1, line 44) is an operation that is independent across compute jobs, and can be performed concurrently. This enables simultaneous transfer of (large) files. Importantly, only file content blobs (i.e., git-annex keys) are transferred at this point.

Additionally, critical metadata must be deposited too. All essential metadata is encoded in the new dataset state commit, recorded on the job-specific Git branch. Consequently, it is deposited using a `git push` call (Listing 1, line 49). This push operation is not concurrency-safe, hence must be protected by a global lock that ensures only one push is performed at a time across all compute jobs (using the tool `flock`). Therefore this step represents a central bottleneck that can influence computational throughput. However, when file-content is only tracked by checksum with git-annex, the changes encoded in the Git branch are metadata only, and a transfer is typically fast.

After successful completion of all computations, the DataLad dataset on permanent storage holds the provenance records of all results in separate job-specific branches, and the content of all output files in a single git-annex object tree.

#### Result consolidation

is the final workflow step. After processing, the result DataLad dataset contains as many branches as successfully completed jobs. These branches must be consolidated into a new state of the mainline branch that jointly represents the outcomes of a individual computations (Figure 1b, “result consolidation/merge”).

How exactly this merging operation must be conducted depends on the nature of the changes. In the simplest case, all compute jobs produced non-intersecting outputs, i.e., no single file was written to by more than one compute job. In this case, all branches can be merged at once using a so-called *octopus-merge*:

~~~
# octopus - merge all “ job “ branches at once
git merge -m “ Merge results” $( git branch - al | grep ‘job -’)
~~~

Depending on the number of result branches, it may be necessary to merge branches in batches to circumvent operating system or shell limits regarding a maximum command line length. If computational outcomes are not independent across jobs (i.e., order of computation/modification matters), a merge strategy has to be employed that appropriately acknowledges such dependencies. If the deposition dataset is hosted in a RIA store (as suggested above for performance reasons) this operation is performed in a temporary clone.

As a final step, valid metadata on output file content availability must be generated. File content resides in the result dataset at the deposition site already, but the required metadata was not pushed to its internal `git-annex` branch from all compute jobs, in order to avoid a consolidation bottleneck. Instead, these metadata are generated only now, by probing the availability of the required file content blobs for all files present in the mainline branch after merging all compute job branches.

~~~
# discover/ confirm result file availability
git annex fsck -- fast -f output - storage
~~~

**Listing 1:**
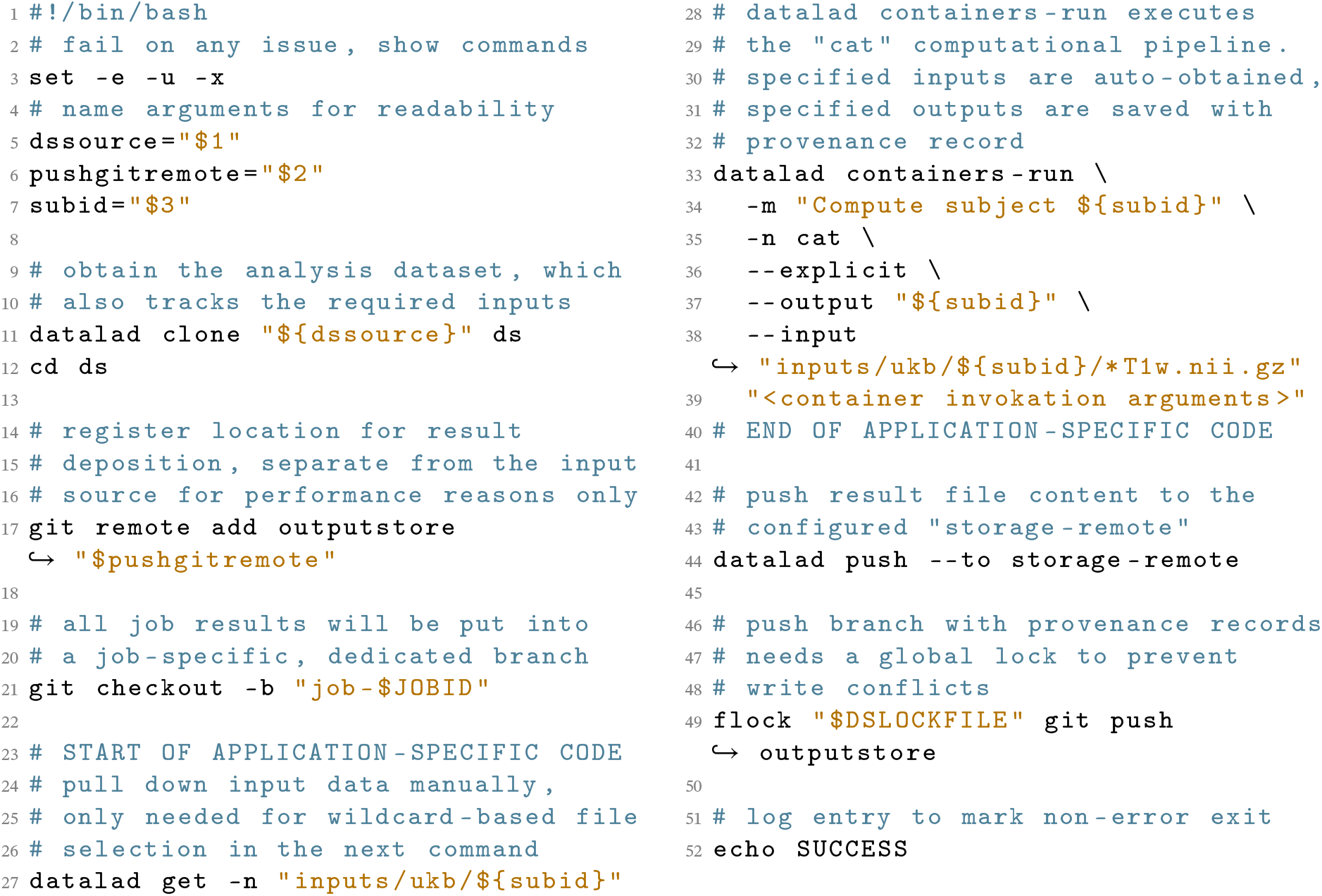
Complete compute job implementation as a bash script. A batch system invokes the job-script in a temporary working directory with three parameters: a URL of a DataLad dataset tracking all code and input data, a URL to deposit job-results at, and an identifier to select a sample for processing. Apart from performance-related optimizations, the job implementation conducts three main steps: 1) `clone` a DataLad dataset with all information to bootstrap an ephemeral computing environment for each job; 2) `containers-run` a containerized pipeline with a comprehensive specification of to-be-retrieved inputs and to-be-captured outputs; 3) push captured outputs and process provenance records to a permanent storage location. Preparation, computation, provenance record creation, and file content deposition on permanent storage are fully independent across jobs, and are executed in parallel. Only the `git push` of the provenance record to a central repository must be protected against concurrent write-access for technical reasons. Additional job parametrization (`DSLOCKFILE` and `JOBID` environment variables) are defined at job-submission using batch system specific means. The job script can be adjusted to a different processing pipeline by replacing the container invocation (see `APPLICATION-SPECIFIC CODE` markers).

~~~
# push consolidated provenance records and file availability
# metadata to permanent storage
datalad push -- data nothing
~~~

The `git-annex fsck` command probes the configured `output-storage` site whether it possesses a given annex key (i.e., a file content blob corresponding to a particular checksum), and generates an appropriate availability metadata record. The final `datalad` push command (Listing 1, line 44) transferred these verified metadata records to permanent storage.

The outcome of this consolidation process is a self-contained DataLad dataset, with valid, machine-actionable provenance information for every single result file of the performed data processing. As such, it is a modular unit of data, suitable as input for further processing and analysis. It translates the advantages of comprehensive and precise linkage of all its components across any number of other data modules to any consumer.

### Balance of reproducibility and performance

Taken together the described approach to reproducible, large-scale computation implements a three layer strategy. From bottom to top, these layers feature different trade-offs regarding portability/reproducibility vs. flexibility for performance adaptations to particular computational environments: The lowest layer is the (`containers-)run` command, comprising an environment specification and instructions to compute the desired outcomes from inputs in this environment. Using suitable technologies, such as computational containers, this layer offers a maximum of portability, but also a minimum of flexibility, as this exact environment must be provided in order to reproduce results. Consequently, the proposed framework captures process provenance at this layer (Figure 2). The middle layer describes how a self-contained, ephemeral workspace can be generated that is suitable for executing the specification of the previous layer (Listing 1). Here, general infrastructural choices can be made. For example, a limitation to a POSIX-compatible environment that is common for HPC/HTC systems, or the granularity with which provenance records are captured (and therefore the granularity at which reproducibility is supported). This layer plays a key role in ensuring that process provenance records are valid and complete. The topmost layer is concerned with maximizing performance on a particular infrastructure via tailored job orchestration, and composition. This layer is poorly portable as it references infrastructure-specific elements, such as job scheduling systems, absolute paths, user names or resource identifiers. While the implementation of all three layers should be provided within the DataLad dataset for a computational project, only the lowest layer is strictly required for reproducing results.

### Software requirements

The software and their minimum version requirements for executing the framework are `datalad` v0.14.2, Git v2.24, `git-annex` v8.202003, and `datalad-container` v1.1.2 (not required for recomputation). Optional requirements are job scheduling systems as well as containerization software (e.g., `Singularity` v2.6.).

In principle, the framework could also be used without a software container. But despite their problems, containers represent the contemporary optimum for encapsulating computational environments that can be shared and reused across different systems. Here we have used Singularity ^15^, one of the most widely used container solutions for both single- and multi-user environments, suitable for HPC/HTC architectures. This choice limits the target platform on which a provenance-based recomputation can be attempted, and for example rules out the Windows operating system for which this software is not available. Other technologies, such as Docker, offer a different set of supported environments.

### UK Biobank computing use case

To demonstrate the framework’s scalability and its ability to create reusable derivatives for subsequent analyses, we applied it to data from the brain imaging component of the UKB project ^16^. We performed a containerized analysis for voxel-based morphometry (VBM) ^28^ based on the Computational Anatomy Toolbox ^29^, a common method for anatomical brain imaging data. This choice of data and processing pipeline posed particular challenges for openness, transparency, and reproducibility. The UKB imaging project is one of the largest studies of this kind. The data underlie strict usage constraints to ensure the responsible use of participants’ personal data. Moreover, the chosen processing pipeline is based on MATLAB, at present still the most prevalent programming environment in biomedical research ^11^, enforcing rigid redistribution limits due to its proprietary, closed-source license. The setup steps were implemented in a bootstrap script available at https://github.com/psychoinformatics-de/fairly-big-processing-workflow/blob/main/bootstrap_ukb_cat.sh.

### Self-contained processing specification as a DataLad dataset

The UKB provides imaging data in ZIP archives, with one archive containing all files for a single modality of a single participant in one format. Direct downloads via versioned perma-URLs are not possible, but `ukbfetch`, a custom binary-only downloader application, must be used.

We implemented `datalad-ukbiobank` ^19^, a DataLad extension (see Box 1) that aids retrieval, indexing, and versioning of UKB data offerings in the form of DataLad datasets. Such a dataset represents data in three variants (using dedicated Git branches): the downloaded ZIP files, extracted ZIP file context using UKB-native filenames, and an alternative data organization following the BIDS standard.

Using `datalad-ukbiobank`, we retrieved MRI bulk data for all participants in NIFTI format. Each participant’s data were represented as an individual DataLad dataset, yielding 42,715 datasets in total. The BIDS-structured branches of all these datasets were jointly tracked by a single UKB superdataset (“Data” in Figure 3). This UKB superdataset is installable within seconds. On a filesystem, it takes up about 40 MB of space, but can retrieve any of the registered file content in the entire DataLad dataset hierarchy, comprising 76 TB across 43 million files, on demand.

As processing pipeline, we chose CAT’s default segmentation of structural T1-weighted images using geodesic shooting ^44^, including calculation of total gray matter (GM), white matter (WM), and intracranial volume (TIV), as well as extraction of regional GM estimates from several brain parcellations. To this end, we built a Singularity container for the MATLAB-based Computational Anatomy Toolbox (CAT; version: CAT12.7-RC2, r1720) ^29^, which is an extension to the Statistical Parametric Mapping software (SPM; version: SPM12, r7771; www.fil.ion.ucl.ac.uk/spm/software). As MATLAB requires a commercial, non-transferable license, we used a compiled version of the CAT toolbox provided by the authors, which does not require the availability of a MATLAB license at runtime. Due to software license restrictions (the MATLAB Compiler Runtime in the container is subject to the MATLAB Runtime license), we cannot redistribute the container image, but we share a detailed description and full recipe of the container together with instructions on how to build and use it at github.com/m-wierzba/cat-container.

We added two custom code files to the dataset. First, a batch script, with a comprehensive specification and parameterization of all processing steps to be performed by CAT in an input image. This script allowed us to bundle up all relevant analysis steps into single command that also defines the smallest unit for recomputation. Second, a utility script to post-process all relevant outputs (≈30 individual files) into four tar archives per computation in order to minimize disk space usage and number of resulting files. Controlling the total number of output files was important due to the amount of computational outcomes to be tracked in this particular result dataset. Only four files per computation translate to more than 160,000 files in total. Such large datasets require substantial file system operations, even when only a subset of file content is retrieved for a particular use case.

The resulting tar archives are organized according to envisioned consumption scenarios (`vbm` containing modulated gray matter density and partial volume estimates in template space, `native` with atlas projections and partial volumes in individual space, `surface` with surface projection and thickness, and `inforoi` containing regional volume and thickness estimates of several atlases/parcellations; Figure 1a). tar was parameterized to create archived with a normalized file order, creation time, and file permissions in order to not introduce artificial variation between recomputations. Likewise, all result files were carefully stripped of timestamps and other non-deterministic log file content. The resulting *reproducible* tarballs allow to attribute file content variability across re-computations to actual result variability.

### Environment and performance optimized orchestration

Data were processed on an HPC cluster and a high-throughput computing (HTC) cluster, each imposing a different set of resource constraints. The HPC system is a modular supercomputer with 1,872 nodes, currently among the 500 fastest compute infrastructures in the world ^45^. While available disk space was abundant, storage was constrained by an inode quota of 4.4 million files – less than the total number of files of the raw dataset. In contrast, the HTC cluster is a mid-sized computational cluster with 31 nodes with only about 400 TB storage capacity, preventing the existence of more than one copy of the raw dataset, and limiting the size of derivatives that could be stored.

To reduce the disk space and inode demands, all DataLad datasets were stored in a RIA store. In this “backend” representation (Figure 1d), a single participant dataset encompasses 25 inodes and about 4 GB of disk space. When cloned into a workspace (Figure 1a), it expands to several hundreds of files. In total, the employed RIA store hosts 42,715 datasets comprising the full UKB data, and consumes 75.6 TB of disk space with less than 940k inodes.

The ability to extract subsets of otherwise compressed inputs only when needed in ephemeral workspaces allowed us to adjust the parallel job load to the available resources. This enabled computations when disk space or inode availability were insufficient for the full dataset. With this setup, we were able to complete data processing for a one-hour-per-image pipeline on the HPC system within 10.5 hours, using 25 dedicated compute nodes, each executing 125 jobs in parallel on RAM disks with GNU Parallel ^46^. On the HTC system in turn, HTCondor scheduled jobs dynamically across several weeks for available compute slots in an otherwise busy system used for unrelated computations by other researchers.

In order to validate different aspects of reproducibility all data processing was performed twice, once on each computing platform, and also a third time for a small subset of the data on a personal laptop. For the two main computing platforms dedicated job submission scripts were implemented for SLURM and HTCondor respectively. In contrast, the partial recomputation on a laptop solely relied on the local availability of the Singularity container technology, but was otherwise fully automatic, based on the captured provenance record, to confirm practical reproducibility for an independent consumer.

Because of the large number of participants in the dataset and the aim to be able to rerun the data processing on a future, even larger release, one compute job per participant was generated. A compute job serially processed either two images, only a single image, or none, depending on the actual data availability. A dedicated provenance record was captured for each pipeline execution on an individual input image, yielding a total of 41,180 records.

### Execution and result consolidation workflow

Data processing was executed based on two variants of the same DataLad dataset, each containing a common computational environment, and the same input data, but a different, optimized job submission implementation. Result consolidation was first performed separately on each computational infrastructure, following the steps described in the framework setup. Lastly, the two complete sets of computational outcomes were integrated in the same dataset, as two different branches, for comparison.

### Result verification

As prior manual data inspection was infeasible due to the amount of data, we included basic checks to ensure availability of T1-weighted images during processing. Subsequent quality control analyses were derived from the computed results. Figure 4 shows the distribution of quality control metrics for T1-weighted images ^47^ across the sample. In addition, we assessed result replicability between recomputations by comparing binary identity of result files between analysis repetitions. To better estimate the amount of dissimilarity between recomputations, we also calculated mean squared error (MSE) over recomputations for a range of key VBM estimates, and the correlations between the brain atlasses included in the CAT toolbox.

### (Re)use

After successful completion, results comprise a collection of different VBM-related measures for all images in the sample, represented in archives. For easier consumption, and as researchers are rarely interested in the full set of measures, the output DataLad dataset was subsampled into smaller “special purpose” datasets. These datasets contained a subset of the results in extracted, and optionally aggregated form, tailored to different research questions, for easier and faster access. This process relied on registering the main result DataLad dataset into a new tailored DataLad dataset via nesting (“Tailored results A/B” in Figure 3), extracting and transforming the required files with provenancetracking by datalad run, i.e., the same mechanism that captured provenance for the initial computation. This approach yields an transparently generated data view that can be updated by re-applying this transformation in case of changed inputs via the `datalad rerun` command.

As a concrete example we generated a DataLad dataset with tissue volume statistics for regions of interests in each parcellation and for all participants. We implemented a script that extracted aggregated noise-to-contrast-ratio, inhomogeneity-to-contrast-ratio, image quality rating, total intracranial volume, total gray matter volume, total white matter volume, total cerebral spinal fluid volume, total white matter hyperintensities volume, and total surface area into one CSV file per brain parcellation. Importantly, we limited the numerical representation of the scores in these tables to an empirically meaningful precision, thereby helping to suppress the undesirable impact of technical side-effects of non-deterministic algorithm implementations and floating point operations on the effective reproducibility of results for any practical purpose. These results are a fraction of the size and number of files of the total results, but sufficient for investigating VBM-related research questions. Using the encoded, machine-actionable provenance information, each result can be traced to the precise files they were generated from in a transparent and reproducible manner.

The direct computational output of the workflow on the UKB sample is therefore not a final result, but an intermediate representation optimized for storage and handling. More tailored views for concrete use cases can be optimized for access convenience. With this, we achieve a compromise between the desires of a data consumer and the demands of the storage infrastructure and operators.

### Open tutorial

As license restrictions prevent open sharing of data and container image used in the UKB showcase, we implemented the processing framework for an additional use case, for which all components can be publicly shared in readily usable form. The resulting, fully populated DataLad dataset is publicly available at github.com/psychoinformatics-de/fairly-big-processing-workflow-tutorial. It can serve as a functional reference implementation that affords reproducibility based on machine-actionable provenance records. All setup steps were implemented in a bootstrap script available at https://github.com/psychoinformatics-de/fairly-big-processing-workflow/blob/main/bootstrap_forrest_fmriprep.sh

#### Self-contained processing specification as a DataLad dataset

As input data we employed a dataset with structural brain imaging data for 20 individuals ^49^ from the studyforrest.org project ^34^, linked as a subdataset at `inputs/data`. This is a BIDS-structured dataset published under the permissive PDDL license. It is publicly available as a DataLad dataset at github.com/psychoinformatics-de/studyforrest-data-structural.

For data processing we use fMRIprep’s structural preprocessing pipeline ^33^ (version v20.2.0) that is freely available as a Singularity container in the DataLad dataset of the public Repronim container registry github.com/repronim/containers. With this pipeline, each T1-weighted MRI scan was corrected for intensity non-uniformity using `N4BiasFieldCorrection` v2.1.0^50^ and skull-stripped using `antsBrainExtraction.sh` v2.1.0 (using the OASIS template). Spatial normalization to the ICBM 152 Nonlinear Asymmetrical template version 2009c ^51^ was performed through nonlinear registration with the `antsRegistration` tool of ANTs v2.1.0^52^, using brain-extracted versions of both T1w volume and template. Brain tissue segmentation of CSF, WM, and GM was performed on the brain-extracted T1w using FAST ^53^ (FSL v5.0.9).

#### Environment and performance optimized orchestration

As both foundational DataLad datasets for input data and pipeline are available from public sources, their file content did not need to be stored on local infrastructure at all. Instead, the processing specification superdataset linked the two components with their GitHub URL, and individual compute jobs retrieved relevant input data from their associated web sources directly.

An example HTCondor-based job-scheduling setup for the HTC infrastructure used in the Open Tutorial showcase is included in the shared resources.

#### Execution and result consolidation workflow

For demonstration purposes the same execution workflow as for the UKB showcase was used. However, due to the small number of compute jobs, and the long individual runtime of each job, implementation details like the separation of clone sources and push targets, or the distinction of result file transfer and provenance metadata deposition only has negligible performance impact.

## Data Availability

Data from the UK Biobank project were obtained from a third party, UK Biobank, upon application. Interested parties can apply for data from UK Biobank directly, at www.ukbiobank.ac.uk.

Structural data from the Studyforrest project ^49^ (doi.org/10.12751/g-node.zdwr8e) are available at github.com/psychoinformatics-de/studyforrest-data-structural. The studyforrest derivatives computed by the tutorial workflow are publicly available from github.com/psychoinformatics-de/fairly-big-processing-workflow-tutorial.

## Code Availability

All scripts used to process the data are publicly available at github.com/psychoinformatics-de/sfairly-big-processing-workflow. The recipe used to build the CAT Singularity container is publicly available at github.com/m-wierzba/cat-container.

## Acknowledgements

The authors wish to thank Timo Dickscheid, Michał Szczepanik and Stephan Heunis for their feedback on earlier versions of this manuscript.

This work was supported by European Union’s Horizon 2020 research and innovation programme under grant agreements Human Brain Project (SGA3, H2020-EU.3.1.5.3, grant no. 945539) and VirtualBrainCloud (H2020-EU.3.1.5.3, grant no. 826421). The development of the DataLad software was supported by grants from the US National Science Foundation (NSF 1912266, 1429999) and the German Federal Ministry of Education and Research (BMBF 01GQ1905, 01GQ1411). MW was supported by ETIUDA grant received from the National Science Centre, Poland (2018/28/T/HS6/00507). This research has been conducted using the UK Biobank Resource under application number 41655.

## Author contributions

ASW, AQW, BP, FH, LKW, MH, and MW conceived the setup of the framework. ASW, LKW, and MH piloted and documented an earlier version of this framework with a smaller dataset. BP, LKW, and MH wrote the software to download and structure UK Biobank to BIDS. FH, MH, and MW containerized the CAT processing toolbox. MH implemented the HTCondor setup and bootstrapping. FH implemented the SLURM setup and conducted QC analysis on the results. ASW, and MW implemented the tutorial on studyforrest.org data. ASW wrote the first draft of the manuscript. ASW, AQW, BP, FH, LKW, MH, MW, and SE contributed to the conceptualization, writing, and editing of the manuscript. All authors read and approved the final draft.

## Competing Interests

The authors declare no competing interests.

## Footnotes

1 A visualization of the different processing speeds can be found at https://www.youtube.com/watch?v=UsW6xN2f2jc.

## Notes

### Competing Interest Statement

The authors have declared no competing interest.

### Summary of Updates

The manuscript was improved with an extended overview of tools sharing similar aims or features in the discussion, among them Parsl, Snakemake, and IPFS, a discussion of the limitations of content-hash-based verification and solutions, and an outlook for adapting the framework to other analyses.

